# Scale-free and oscillatory spectral measures of sleep stages in humans

**DOI:** 10.1101/2022.06.09.495470

**Authors:** Bence Schneider, Orsolya Szalárdy, Péter P. Ujma, Péter Simor, Ferenc Gombos, Ilona Kovács, Martin Dresler, Róbert Bódizs

**Author notes:** Correspondence: Bence Schneider.

## Abstract

Power spectra of sleep electroencephalograms (EEG) comprise two main components: a decaying power-law corresponding to the aperiodic neural background activity, and spectral peaks present due to neural oscillations.’Traditional’ band-based spectral methods ignore this fundamental structure of the EEG spectra and thus are susceptible to misrepresenting the underlying phenomena. A fitting method that attempts to separate and parameterize the aperiodic and periodic spectral components called ‘fitting oscillations & one over f’ (FOOOF) was applied to a set of annotated whole-night sleep EEG recordings of 251 subjects from a wide age range (4-69 years). Most of the extracted parameters exhibited sleep stage sensitivity; significant main effects and interactions of sleep stage, age, sex, and brain region were found. The spectral slope (describing the steepness of the aperiodic component) showed especially large and consistent variability between sleep stages (and low variability between subjects), making it a candidate indicator of sleep states. The limitations and arisen problems of the FOOOF method are also discussed, possible solutions for some of them are suggested.

## 1 INTRODUCTION

The observation that besides the oscillatory activity of the brain there is an aperiodic background component that manifests itself as a power-law in the power spectra of electroencephalography (EEG) signals is not a novelty [Matthis et al., 1981; Pritchard, 1992], however the approach to characterize the whole spectrum with only a small number of parameters that describe the periodic and aperiodic spectral components separately is becoming increasingly relevant [Donoghue et al., 2020; Bódizs et al., 2021b].

Traditional methods in EEG analysis often define fixed frequency bands and then search for differences in the spectral power of these. The bands are supposed to correspond to neural oscillations, however testing the sum power of a fixed frequency band might be misleading as the emerging effect could indeed reflect the changes in the power of the oscillatory activity, but also a shift in the oscillation frequency or a change in the aperiodic background activity. Another drawback of the band-based methods is that the spectral power of the bands can be significantly correlated (due to the fact that in lack of oscillations they all represent a portion of the same overarching power-law) and thus carry redundant information. By the independent parametrization of the aperiodic and periodic power spectrum components both of these problems are eliminated.

At the same time the significance of the so-called ‘spectral slope’ is also gaining focus in recent studies in the field of electrophysiology and neuroscience. The power-law in the power spectrum can be described in the form: *P*(*f*) ∝1/*f^s^*, *s* > 0 or (where *P* is the power and *f* is the frequency, and the exponent *s*), equivalently it can be written as *P* (*f*) ∝ *f^x^* in which case the exponent for the decaying case is negative *x* = −*s*, plotting this type of relationship on a double logarithmic scale results in a linear function with the slope being equal to the power exponent x. Signals that typically present such scale-free power spectra are ‘colored’ or ‘1/f noises’, most famous mathematical examples being the white noise (*x* = 0), pink noise (*x* = −1), and brown noise (*x* = −2). Real-world examples include shot-noise in electronic devices, but the 1/f spectrum has been discovered on many different levels in the nervous system and the brain: in the fluctuations of membrane potentials, in local field potentials (LFP) [Baranauskas et al., 2012], electrocorticography (ECoG)[Zempel et al., 2012] and in electro- and magnetoencephalography signals (EEG, MEG) [Bénar et al., 2019]. Furthermore, the spectral slope exhibits physiologically and medically relevant effects: it had been found to change with aging [Voytek et al., 2015], proved to be a significant marker of schizophrenia [Racz et al., 2021] and attention deficit hyperactivity disorder [Karalunas et al.,2022], and also an indicator of consciousness during anesthesia [Colombo et al., 2019].

The most conspicouos physiological changes in brain electrodynamics appear during the changes in wake-sleep states [Lázár AS, 2022]. Although there are a few reports on sleep-related changes in scale-free, aperiodic EEG activity [Bódizs et al., 2021b; Miskovic et al., 2018; Lendner et al., 2020], a comprehensive depiction of sleep stage-dependent variation in all scale-free and oscillatory parameters is still lacking. In the present study we describe the oscillatory and scale-free spectral parameters of sleep stages by applying a fitting method to the power spectra of a large EEG data set and show that many of these parameters show sleep stage, age and sex effects and interactions. Moreover, after correcting for individual differences the spectral slope proves to be an effective indicator of sleep stages.

## 2 MATERIALS AND EQUIPMENT

The Budapest-Munich database of sleep records contains whole night EEG/polygraphy signals derived from 251 healthy subjects, 122 females in the age range of 4-69 years, and were divided into age-groups of children (4-10 years, N = 31), teenagers (10-20 years, N = 36), young adults (20-40 years, N = 150) and middle-aged adults (40-69 years, N = 34) [Bódizs et al., 2021a]. The standard 10-20 electrode placement was used, out of which 10 channels were used in this study (Fp1, Fp2, F3, F4, P3, P4, C3, C4, O1, O2), see Figure 1. The database is a collection of records performed in multiple laboratories with different sampling rates, precisions and filter settings (Table 1), but common in terms of covering a whole night of undisturbed sleep following an adaptation night (the latter was not used in the present study). EEG records were offline re-referenced to the mathematically-linked mastoids before being subjected to quantitative analyses. The recordings were scored in 20 s according to the American Academy of Sleep Medicine (AASM)2 coding rules and artefact annotated 4 s segments [Berry, 2018].

**Figure 1.**
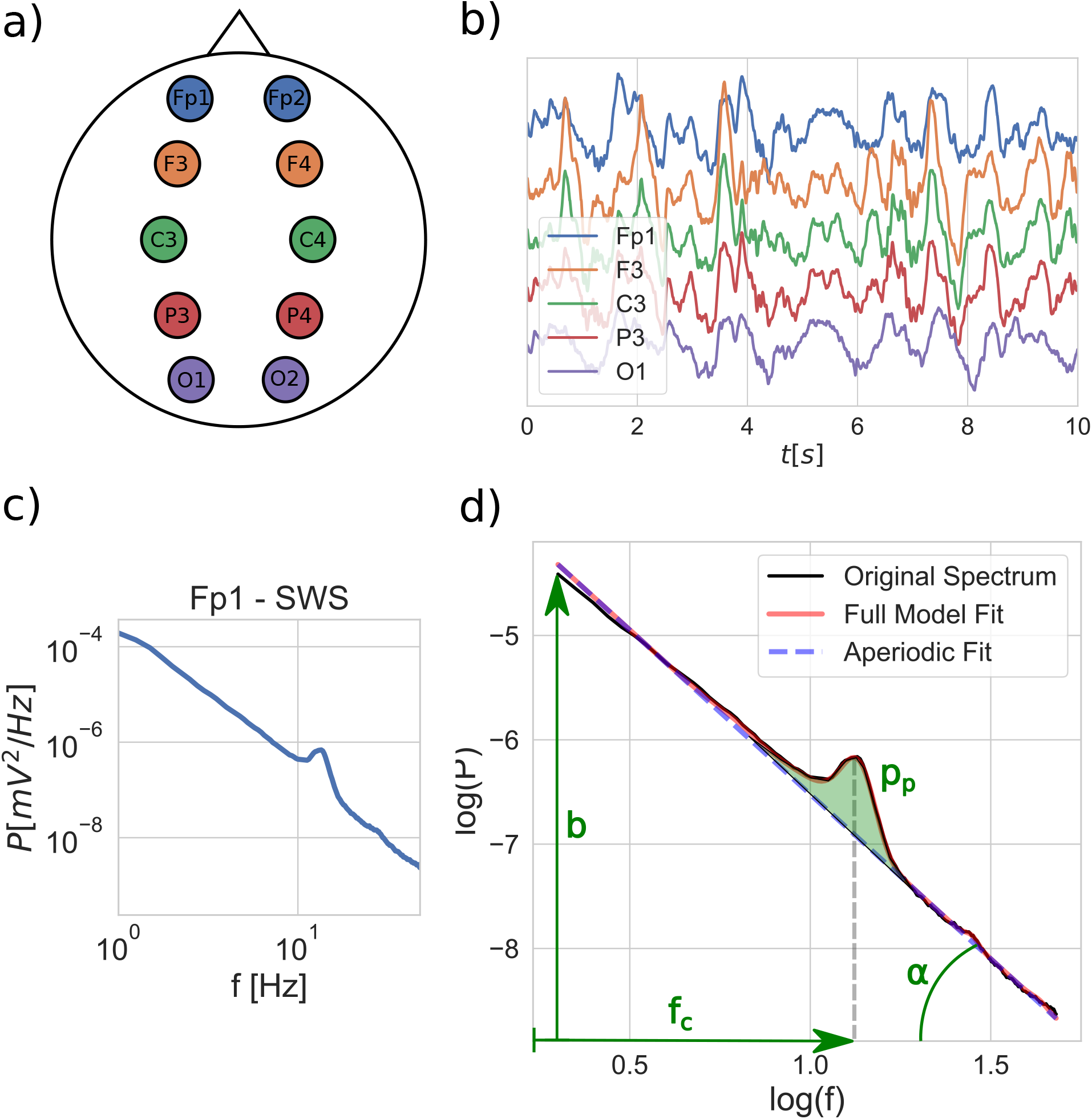
Schematic outline of the fitting process: a) EEG is recorded with electrodes placed according to the 10-20 system, out of which 10 are used, covering 5 brain regions on both left and right hemispheres. b) Time domain signals are segmented into 20-second windows and grouped by sleep stages for each EEG channel. c) Average power-spectral density is calculated for each sleep stage per channel. d) The FOOOF model is fitted to the average power spectra, and the model parameters are extracted: spectral slope (*x* = *tan*(*α*)), intercept (*b*), peak central frequency (*f_c_*) and peak power (*p_p_*).

## 3 METHODS

### 3.1 Power spectrum calculation

The signals of every EEG channel had been segmented by a 4 second sliding window with a 2 second step (50% overlap). Windows that included artifacts were ignored. Each window was Hanning-tapered before their FFT was computed by the use of a mixed-radix procedure. For every subject and channel the segments were grouped by sleep stage, then Welch’s method was applied to obtain the average power-spectral density.

### 3.2 Model fitting

The ‘fitting oscillations & one over f’ (FOOOF) method was used to extract the parameters of the spectra, namely: the spectral slope and intercept and for each spectral peak their central frequency, power and bandwidth. The FOOOF method introduces a physiologically-informed model that attempts to describe neural power spectra (eq.1) as the compound of a power-law representing the aperiodic component (eq. 2), and any number of Gaussian functions approximating the oscillatory peaks (eq. 3).

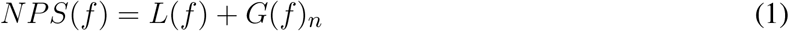

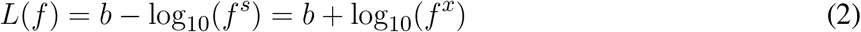

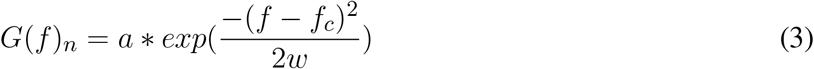

The method calculates a first approximation of the aperiodic components and subtracts the power-law from the original spectrum to achieve a flattened version of the spectrum exposing the spectral peaks. In the next phase in order to find the periodic components Gaussians are iteratively fitted and subtracted from the flattened spectrum. Finally the total periodic component is removed from the original spectrum resulting in a peak-removed spectrum and the aperiodic component is fitted again. The final result is the combination of the aperiodic and periodic components.

After testing the FOOOF method on a subset of our data, it could be observed that using the default setting values provided by the method leaded to over- or underfitted results in multiple cases (see details in supplement). After looking at the problematic cases individually and assessing the underlying causes for the faulty fits we found a combination of setting values that yielded adequate results (goodness of fit R-squared values min.: 0.6401, mean: 0.9908, max.: 0.9999). The fitting range was set to the 2-48 Hz frequency interval, the bandwidth of the accepted peaks to the 0.7-4 Hz range and the peak threshold lowered to 1.

### 3.3 Parameter extraction and statistical analysis

After having all the fitted parameters a number of them were selected for analysis: the spectral slope, the center frequency and power of the spectral peak that had the highest power. As the spectral intercept calculated by the FOOOF method is heavily correlated with the slope and doesn’t provide substantially more information, we included in our analysis an alternative spectral intercept also described here ([hivatokozas]) that is defined as the y-axis intersection of the fitted power-law at the frequency location of the largest oscillatory peak (see supplementary information for more details).

General linear model analysis (repeated measures ANOVA with sigma restricted parametrization) had been carried out for each extracted spectral parameter as the dependent variable with the categorical factors of sex (female and male) and age group (child, teenager, young adult, middle aged adult). The within-effects considered were sleep stage (wake (AASM stage: W), non-rapid eye movement sleep 1 & 2 (NREM1 and NREM2 indicating the AASM categories of N1 and N2, respectively), slow-wave sleep (SWS, AASM category: N3) and rapid-eye movement sleep (REM, AASM stage: R), brain region (frontopolar, frontal, central, parietal, occipital) and laterality (left and right). Several main and interaction effects were found, in the following subsections we highlight the most significant ones, for the complete statistical report see the supplementary material.

## 4 RESULTS

### 4.1 Spectral slope

EEG spectral slopes strongly depended on sleep stages (*F*(4, 824) = 770.29, *p* < .00001, 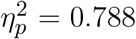) (see Figure 2.). The average slope value was highest in the wake state and decreased as the sleep deepened through the NREM sleep stages, reaching its lowest value during SWS. Also the main effect of age was significant indicating steeper slopes in younger sunjects (*F*(3, 206) = 6.47, *p* < .0001, 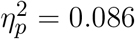). Reliable topographical differences in spectral slopes were evidenced by the main effect of brain region (*F*(4, 824) = 113.33, *p* < .00001, 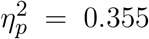). The latter findings indicate steeper slopes in more anterior recording sites. The most significant interactions were the stage-region (*F*(16, 3296) = 55.23, *p* < .00001, 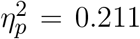) and the stage-region-age interaction (*F*(48, 32) = 4.95, *p* < .00001, 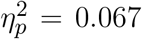), see Figure 2., where the age effect is also depicted. The spectral slope increases with age, reflecting a shallower sleep especially in the SWS stage, being consistent with the known phenomenon of sleep quality deterioration of middle-aged and older adults.

**Figure 2.**
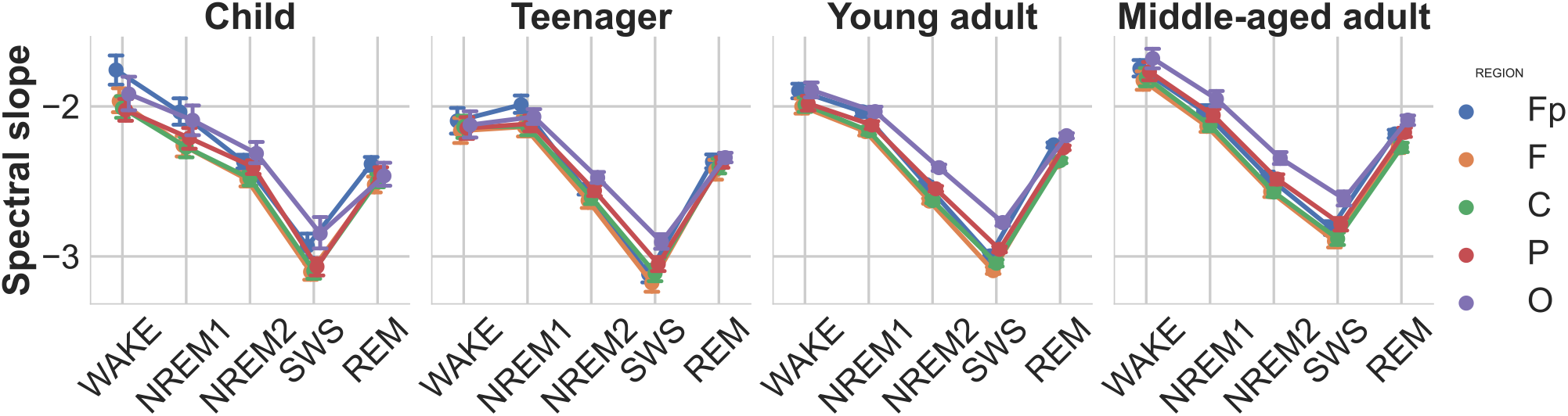
Spectral slopes as functions of sleep stage, brain region, and age group. Note the gradual decrease of slope values (decreasing spectral exponents, increasing steepness) during the course of deepening of NREM sleep, as well as a relatively increased slope in REM sleep (but still below the NREM1 values). Vertical bars denote 95% confidence intervals.

### 4.2 Intercept

As the intercept parameter provided by the FOOOF method is correlated to the slope parameter, we adopted an alternative intercept measure calculated at the frequency of the largest peak, for which we also found a sleep stage main effect (*F*(4, 836) = 35.73, *p* < .00001, 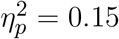) indicating increased intercepts in the slow-wave sleep stages, an age main effect (*F*(3, 209) = 26.37, 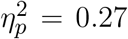) the intercept being higher in children, furthermore a stage-region interaction (*F*(16, 3344) = 9.18, *p* < .00001, 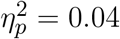, see Figure 4) showing the increase being more pronounced in the frontopolar and frontal regions.

**Figure 3.**
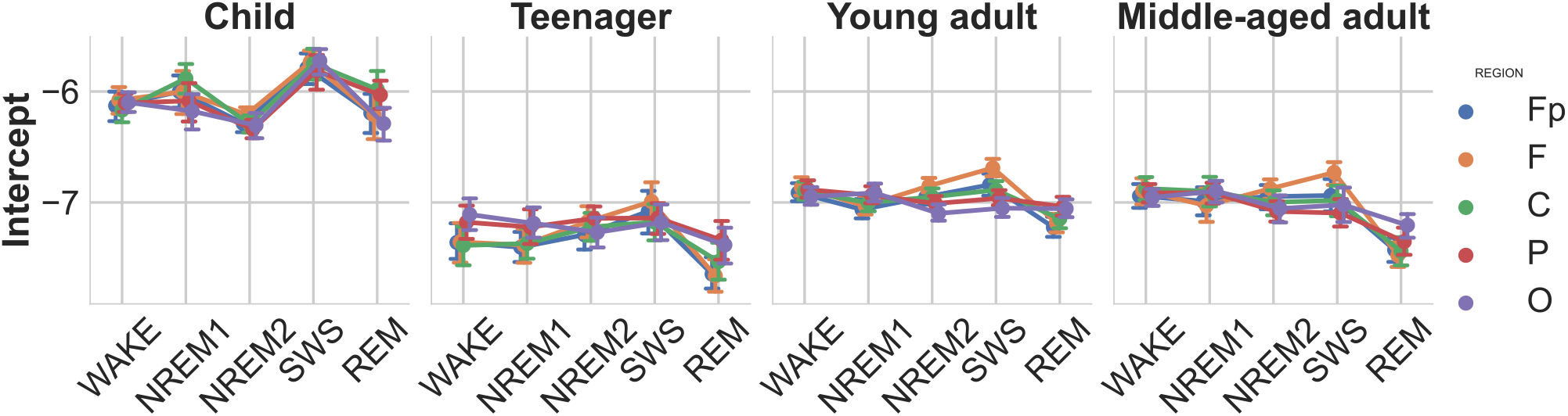
Modified spectral intercepts as functions of sleep stage, brain region, and age group. Note the particularly high intercepts in children, indicating high overall EEG amplitude values. Furthermore, the modified intercepts of SWS stage, especially the ones measured over the frontopolar recording regions, exceed other stages and regions. Vertical bars denote 95% confidence intervals.

**Figure 4.**
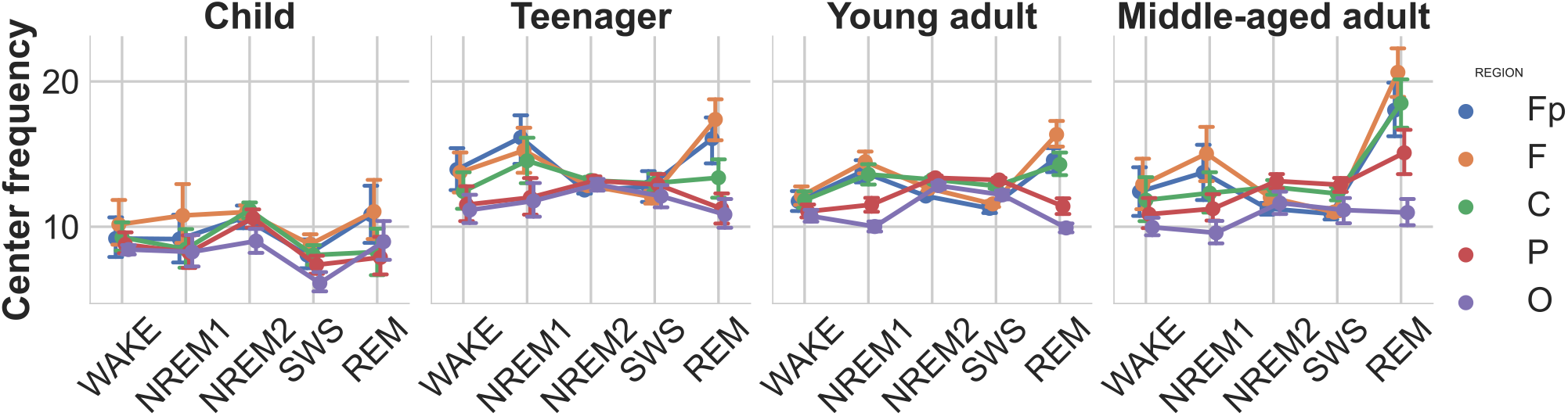
Central peak frequencies as functions of sleep stage, brain region, and age group. Note the high intersubject variability of central peak frequencies in WAKE, NREM1 and REM stages, as compared to NREM2 and SWS frequencies. This pattern indicates the presence of multiple oscillators with individually variable dominance in WAKE, NREM1 and REM stages, as well as a reliable dominance of sleep spindle waves (11-16 Hz) in NREM2 and SWS. Vertical bars denote 95% confidence intervals.

### 4.3 Peak central frequency

The central frequency of the peak with the highest power was more increased in the frontal and frontopolar regions (main effect of region: *F*(4, 824) = 58.138, *p* < .00001, 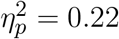). There were main effects of age (*F*(3, 206) = 17.2,*p* < .00001, 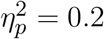) and sleep stage (*F*(4, 824) = 15.584, *p* < .00001, 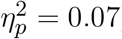) as well. It can be observed that the dominant peak frequencies converge to the characteristic sleep spindle frequency in the NREM2 stage, becoming the most consistent in teenagers.

### 4.4 Peak power

Similarly to the previous parameters, significant main effects of sleep stage (F(4, 824) = 88.765, *p* < .00001, 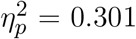) and brain region (F(4, 824) = 97.645, *p* < .00001, 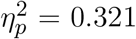) were found for the power of the strongest spectral peak as well. Stage-region and stage-region-age interactions also occurred. On Figure 5 it is interesting to note how the power of the spectral peak in the NREM2 stage becomes prominent in teenagers/young adults and then declines with aging.

**Figure 5.**
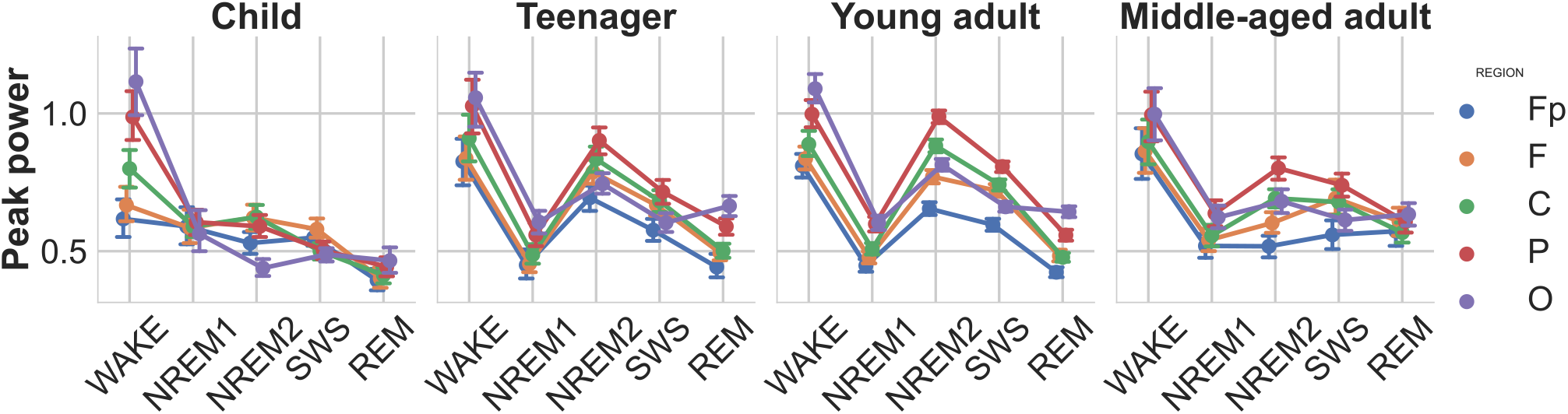
Power of the largest spectral peak as function of sleep stage, brain region, and age group. Note the high peak power in wakefulness and NREM2 sleep, known to be characterized by prominent alpha and sleep spindle oscillations, respectively. In addition, peak power is lower in children and in middle aged adults, as compared to teenagers and young adults. This pattern coheres with the ontogeny of sleep spindle oscillation in humans. Vertical bars denote 95% confidence intervals.

### 4.5 Adjusted spectral slope

Recent findings suggest that the spectral slope is subject-specific, and characterized by high individual-specificity and repeatability [ref]. Furthermore, we found that the slopes of sleep stages are also significantly correlated within subjects. In an attempt to remove this specificity and obtain a subject-independent measure that reflects the sleep stages even more clearly, we introduce the adjusted spectral slope, which takes the slope value of the wake state as the individual reference value by subtracting the slope of the wake stage from all other stages of the subject. In other words the adjusted spectral slope is the deviation from the baseline slope of the wake stage. This new measure showed an even stronger sleep stage main effect (*F*(3, 627) = 1198.56, *p* < .00001, 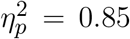), and also a stage-region interaction (*F*(12, 2508) = 62.99, *p* < .00001, 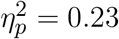).

## 5 DISCUSSION

The application of quantitative EEG methods are widespread and have a long history in the field of sleep research [Cox and Fell, 2020], however there are core aspects of sleep that still lack proper quantifiability, (eg. the scoring of EEG recordings is done (semi-)manually, precisely because no reliable objective measure yet exists that is comparable between subjects and from which a hypnogram could be derived directly). In the present study we have found that the parameters fitted to the spectra depended on the sleep stage and in many cases a large amount of their variance could be explained by the effect of sleep stage, indicating that the parametric description of the EEG power spectrum which separates the scale-free and oscillatory components is also meaningful when describing sleep states. The spectral slope proved to be an especially strong indicator of sleep stages, its consistency improved further by the suggested adjustment (i.e. using wakefulness as a normalizing factor). The nominal values of the EEG spectral slopes suggest that the overall group mean specific to wakefulness is slightly above −2, whereas sleep per se is below this border. The spectral slope equalling the value of −2 is indicative of Brownian motion. Above and below this value the antipersistent and the persistent fractional Brownian motion are found, respectively. That is, our findings suggest that the monopolar, linked-mastoid referred EEG is characterized by antipersistent Brownian motion during wakefulness (with successive increments characterized by overall negative correlation), whereas sleep is best described by persistent Brownian motion (successive increments correlating positively). Former studies focusing on the fractality and smoothness of the EEG signal as expressed in terms of the Hurst-exponent reported similar findings [Weiss et al., 2011].

Although a recent report revealed the individual-specificity of EEG spectral slope values [McSweeney et al., 2021], no former study focused on the between state consistency of this effect. Here we report reliable positive correlations between spectral slopes assessed in different sleep stages, which we consider an important aspect of the individual fingerprint-aspect of brain electrodynamics. Given the finding that within-subject consistency of EEG spectral slopes transcend sleep stages, we performed an adjustment of state-specific slope values by normalizing them against resting wakefulness-derived values. As regarding this adjusted deepening of sleep expressed by changes in the spectral slope relative to wakefulness, our findings are even more reliable than the outcomes based on the absolute values of spectral slopes. Sleep stages can be characterized by a fine-tuned decrease of the spectral exponent relative to the wake state, the decrease ranging from −0.2 to −1 in the states of NREM1 and SWS, respectively. The exceptionally high effect size characterizing this stage-dependency suggests particularly reliable differences and a predictable sequence of changes during the sleep process. Such findings suggest that well-fitted spectral slopes are ideally suited to be the basis of an automated sleep analysis and sleep staging procedure.

In addition to revealing sleep stage effects, our current findings confirm the age- and region-dependency of scalp-recorded EEG spectral slopes reported by former studies [Voytek et al., 2015; Bódizs et al., 2021b; Pathania et al., 2022]. Spectra is steeper in younger subjects and in more anterior recording locations. The first order interactions reported in our current study suggest that the effects of age and region on spectral steepness are unevenly distributed over different sleep stages. Most conspicuous age-related EEG spectral slope flattening is evident in SWS. Likewise, antero-posterior differences in EEG spectral steepness are particularly prominent in SWS. Moreover, these regional differences vary as a function of age (significant region × age group interaction) indicating relatively lower antero-posterior differences in EEG spectral steepness of children and teenagers as compared to adults, as well as unusually steep spectral slopes in the wake state of teenagers.

Intercepts of the spectra in the log-log plane were shown to strongly reflect spectral slopes, with higher intercepts reflecting steeper slopes. This interdependence was hypothesized to reflect the phenomenon of non-zero intercepts, meaning that the spectral slopes are revolving around specific frequencies not equalling ln1 = 0. In order to detect the location of these slope-independent intercepts, we run a series of correlational analyses in our former study, revealing that NREM sleep EEG spindle frequencies (ln12.2 and ln13.5) are ideal candidates for these points, as they resulted in intercepts which are statistically unconnected the spectral steepness (Bódizs et al., 2021). Given the unequivocality of spectral peak emergence in the spindle range of NREM sleep EEG, we intended to determine the spectral intercepts at the center peak frequency in the current study (assuming that slope-independency of spectral intercepts can be found under this peaked sections of the spectra). Our findings indicate the independence of this alternative and adaptive spectral intercept from the slope, which contrasts the correlations of classical intercepts and slopes. That is, we can provide non-redundant sleep stage-effects when analyzing these alternative spectral intercepts.

Central peak frequencies were assumed to reflect the neural oscillatory peculiarities of different sleep stages, indicating specific oscillatory mechanisms known to be operative in specific behavioral states: alpha activity (8-12 Hz) in resting wakefulness, sleep spindle frequency activity (11-16 Hz) in NREM sleep stages and perhaps SWS as well, and theta (4-8 Hz) or beta (16-30 Hz) in REM sleep. In addition, we tested if FOOOF is instrumental in differentiating slower anterior sleep spindle oscillations from faster, more posterior ones. Our initial findings lead us to readjust the standard settings of the FOOOF procedure in order to avoid the amalgamation of two adjacent spectral peaks into a single one, with broader frequency dispersion. After this correction we obtained significant sleep stage and region effects, as well as an interaction of these two factors. Thus, findings indicate reasonable state and localizational effects in oscillatory EEG frequencies. However, the nominal frequencies in the wake state vary as a function of age and provide a stable alpha frequency in children only. In turn, NREM 2 and SWS sleep stages are characterized by prominent sleep spindle frequencies, with detectable antero-posterior differences. NREM1 and REM sleep are characterized by beta oscillatory frequencies, with striking antero-posterior differences: in contrast with NREM2 and SWS faster oscillations are peculiar to anterior sites in these states. The similarities of NREM1 and REM sleep EEG spectra were reported earlier [Bódizs et al., 2008]. Our current findings indicate that besides band-limited power values, the similarity of NREM1 and REM sleep stages is evident in terms of the regional distribution of oscillatory peak frequencies as well.

High peak power values were found to be characteristic features of wakefulness and NREM2 sleep, lowest values in NREM1 and REM, as well as intermediate ones in SWS. These findings fit the knowledge on the prominent alpha and sleep spindle oscillations in wakefulness and NREM2 sleep, respectively. Sleep spindles were also termed as hallmarks of stage 2 sleep. This assertion coheres well with our current findings. Age-related changes in peak power indicate a biphasic change in NREM2 sleep spindle frequencies: initial increase peaking in teenage/young adult years, followed by a decrease in middle aged adults. Again this finding coheres well with reported age-dependent changes in sleep spindle activity [Purcell et al., 2017].

Based on the above findings we conclude that the spectral parameters derived from the fine-grained differentiation of scale-free and oscillatory activities of the EEG are potentially suited to serve as objective measures characterizing sleep states paving the way toward an automatic evaluation of the process of human sleep. Future investigations have to reveal the potential computational and physiological relevance of parametrizing aperiodic and oscillatory activity during the course of the human sleep-wake cycle (i.e. by transcending epoch-based expert scoring). We consider the current findings as a promising first step toward an automatic and objective characterization of sleep-wake dynamics.

## 6 ACKNOWLEDGMENTS

Research supported by the Hungarian National Research, Development and Innovation Office (K-128117; https://nkfih.gov.hu/about-the-office), the Ministry of Innovation and Technology of Hungary from the National Research, Development and Innovation Fund, financed under the TKP2021-EGA-25 funding scheme, the Netherlands Organization for Scientific Research (NWO; https://www.nwo.nl/en), the European Cooperation in Science and Technology (COST Action CA18106; https://www.cost.eu/), as well as the general budgets of the Institute of Behavioural Sciences, Semmelweis University (http://semmelweis.hu/magtud/en/) and the Max Planck Institute of Psychiatry (https://www.psych.mpg.de/en). The funders had no role in study design, data collection and analysis, decision to publish, or preparation of the manuscript.

## 7 SUPPLEMENTARY MATERIALS

### 7.1 Alternative intercept rationale and definition

One of the biggest advantages of adopting a parametric model for describing EEG power spectra is that we can capture the spectral phenomena using significantly fewer variables compared to the original spectral data (2 aperiodic parameters + 3 per peak, whereas the minimal number of bins in the power spectra is generally around 256), however we found that the slope and the intercept parameters provided by the FOOOF method are correlated with an average correlation of: < *r* >= 0.47 (Pearson correlations were calculated between the two variables for all the corresponding sleep stages and EEG channels, then averaged in the Fisher z-space and inverse transformed). The alternative intercept was defined as the value of the power-law component at the frequency of the peak with the highest power, as suggested in an earlier study, in order to achieve the least correlation with the slope. The average correlation between slope and the alternative intercept was < *r* >= −0.03.

### 7.2 Between-stage correlations in the spectral slope

The strongly subject-specific nature of the spectral slope had been demonstrated before, however only in the wake, resting state. In order to test this specificity in the domain of sleep, we compared spectral slopes values between sleep stages and found that in general there is a positive correlation between all stages within individuals, furthermore that the correlation is stronger between subsequent sleep stages, see Figure 7. Correlations coefficients between non-identical stages were in the range of 0.2 < *r* < 0.8, with overall average correlation of < *r* >= 0.49, while p-values were between 10^-58^ and 10^-2^.

**Figure 6.**
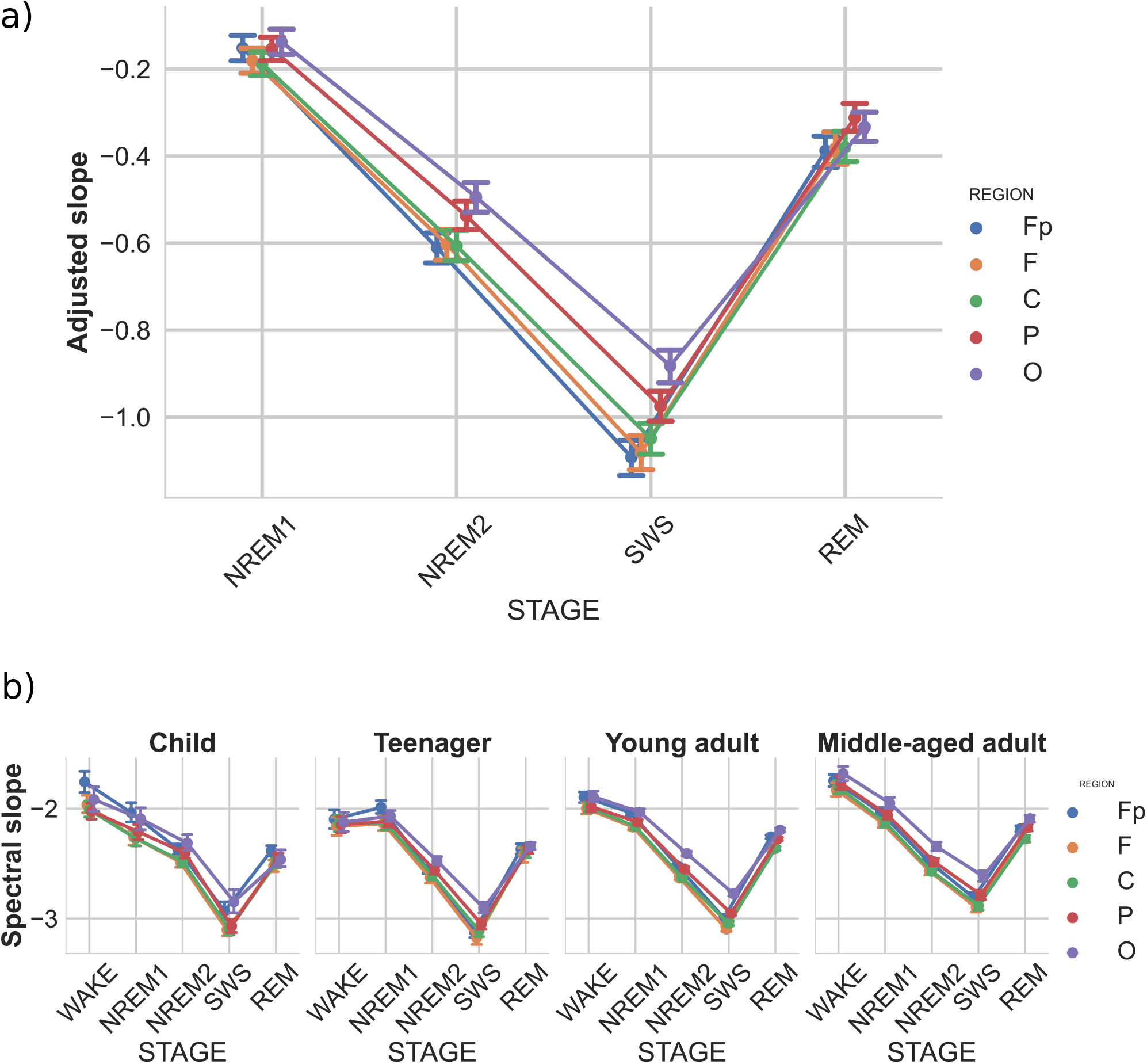
Adjusted spectral slopes as functions of sleep stage, brain region, and age group. Slopes were expressed as deviations from individual-, sleep stage- and recording location-specific deviations from corresponding resting wakefulness values. a) Overall group mean. b) age-group-specific means. Note the particularly reliable steepening of spectral slopes from NREM1 through NREM2 to SWS (indicated by decreasing slope values), as well as a considerable flattening in REM sleep, slightly above NREM2, but below NREM1-specific values. Vertical bars denote 95% confidence intervals.

**Figure 7.**
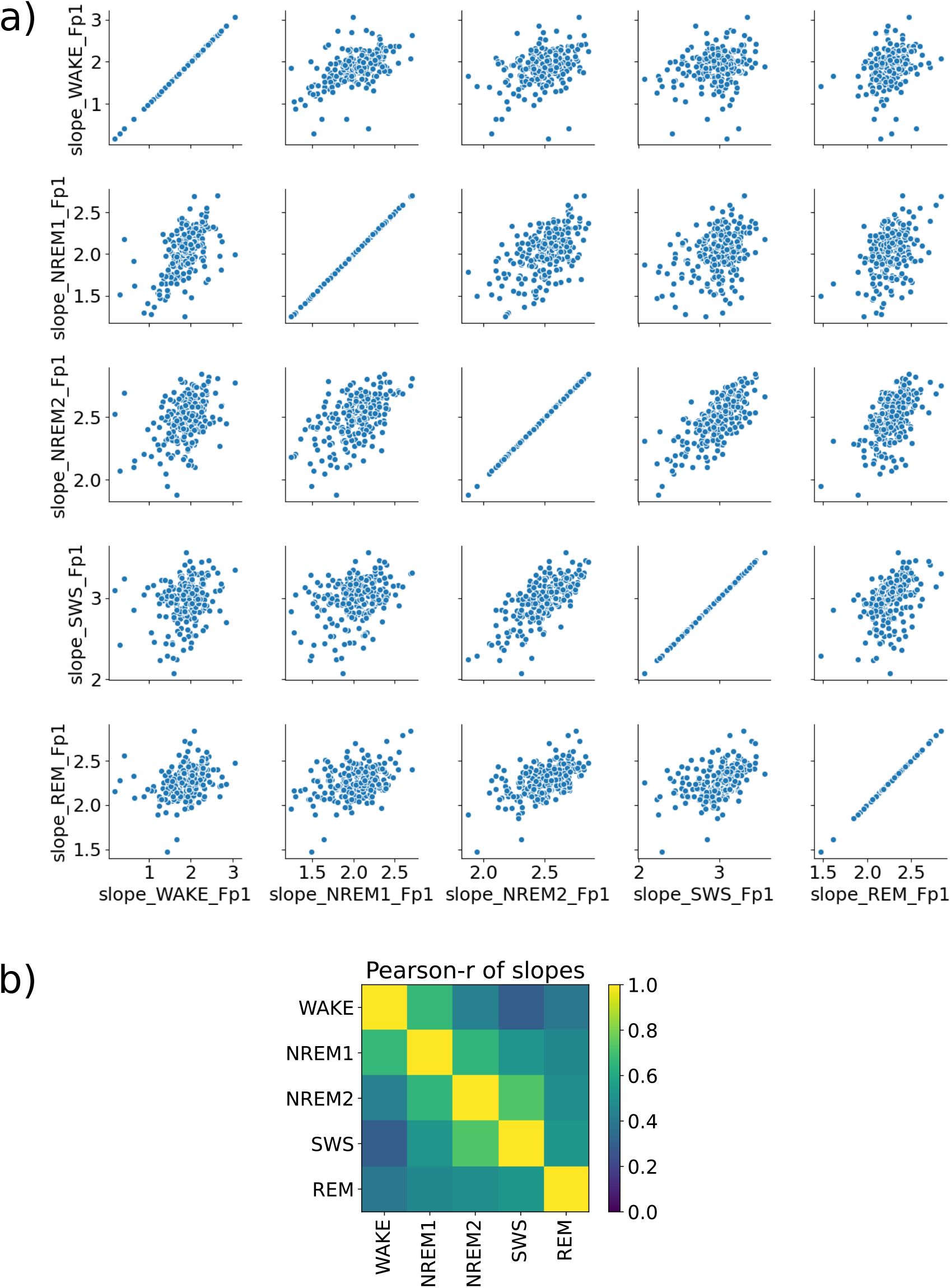
a) Covariance plot of spectral slopes between different sleep stages, each point corresponding to a subject. b) Pearson correlations of spectral slopes between sleep stages in the case of the Fp1 channel. It can be noted that the values closer to the main diagonal are higher, suggesting that there is more correlation between subsequent sleep stages.

### 7.3 Challenges using the FOOOF method

Applying the FOOOF method to spectral data is straightforward, however by looking more closely at certain cases some undesired effects were noticed regarding the periodic component. In some instances when power spectra contained ‘double’ peaks, the model fitted them wrongly as a single wide peak (see below on Figure 8 a) subfigure’s lower row). In order to eliminate this issue the peak threshold parameter was decreased, which ultimately resulted in an acceptable fit, yet it was also an indication that more careful choice of control parameters might be needed than primarily expected. (The peak threshold determines the minimal deviation in power from the aperiodic component that is necessary for the data point to be considered as a peak candidate.) In order to investigate the effect of the peak threshold the same power spectrum had been fitted multiple times while varying the peak threshold quasi-continuously. On Figure 8 b) the frequencies of the found peaks were plotted in function of the peak threshold parameter, it had been expected that the number of found peaks increases as the threshold value descreases (as smaller irregularities in the power spectrum have a higher chance to be above this threshold), however a frequncy shift was also discovered in function of the peak threshold parameter. Knowing that the central peak frequencies are sensitive to this parameter choice a supervision of the fitting is advised.

**Figure 8.**
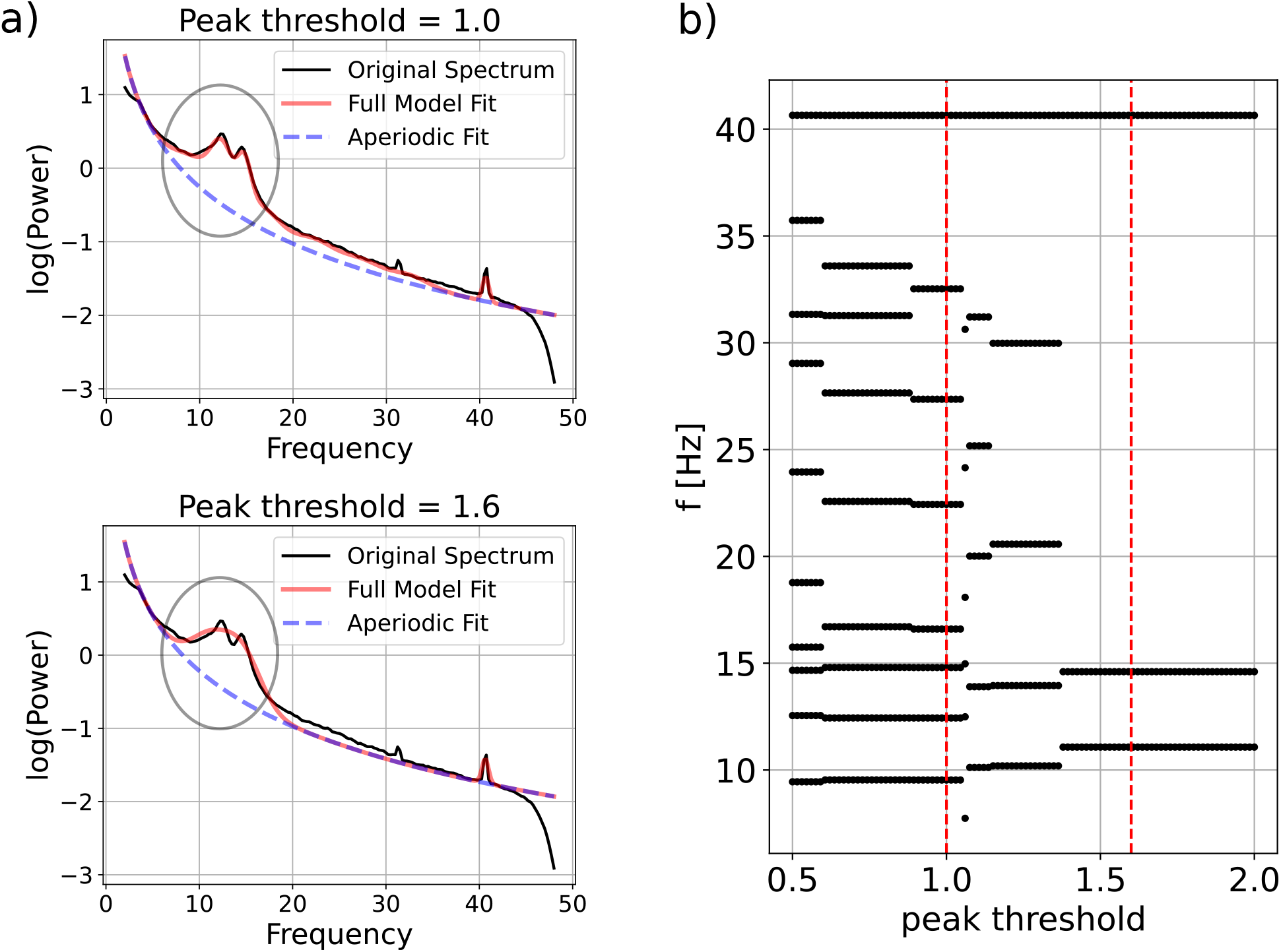
a) Two examples of fitting the same power spectrum using the FOOOF method with different peak threshold parameters, the model detects the double peak only for lower peak thresholds. b) Dependence of fitted peak frequencies on the peak threshold parameter. The vertical dashed lines represent the two cases on the left.

### 7.4 Statistical results

#### 7.4.1 Spectral slope

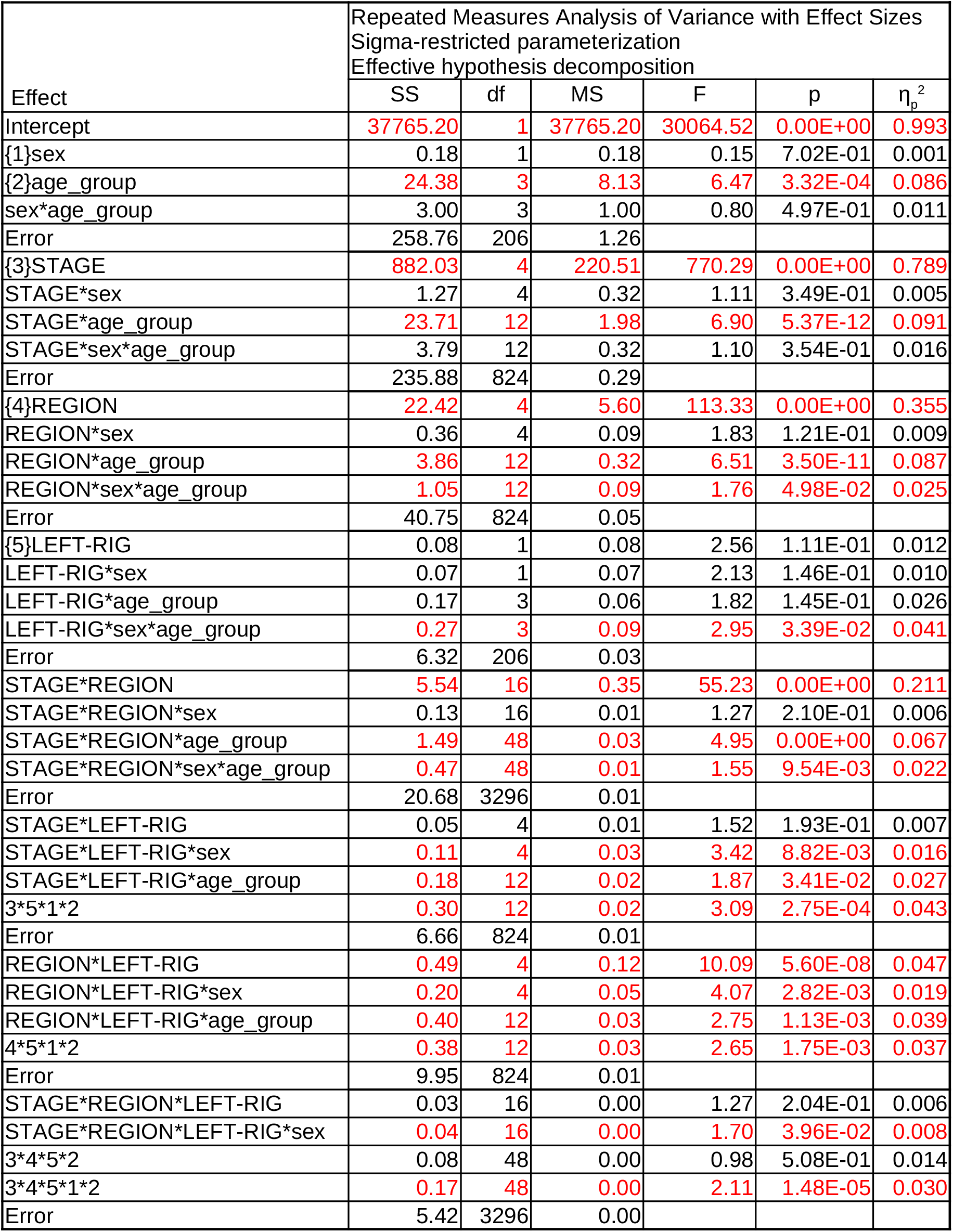

#### 7.4.2 Intercept

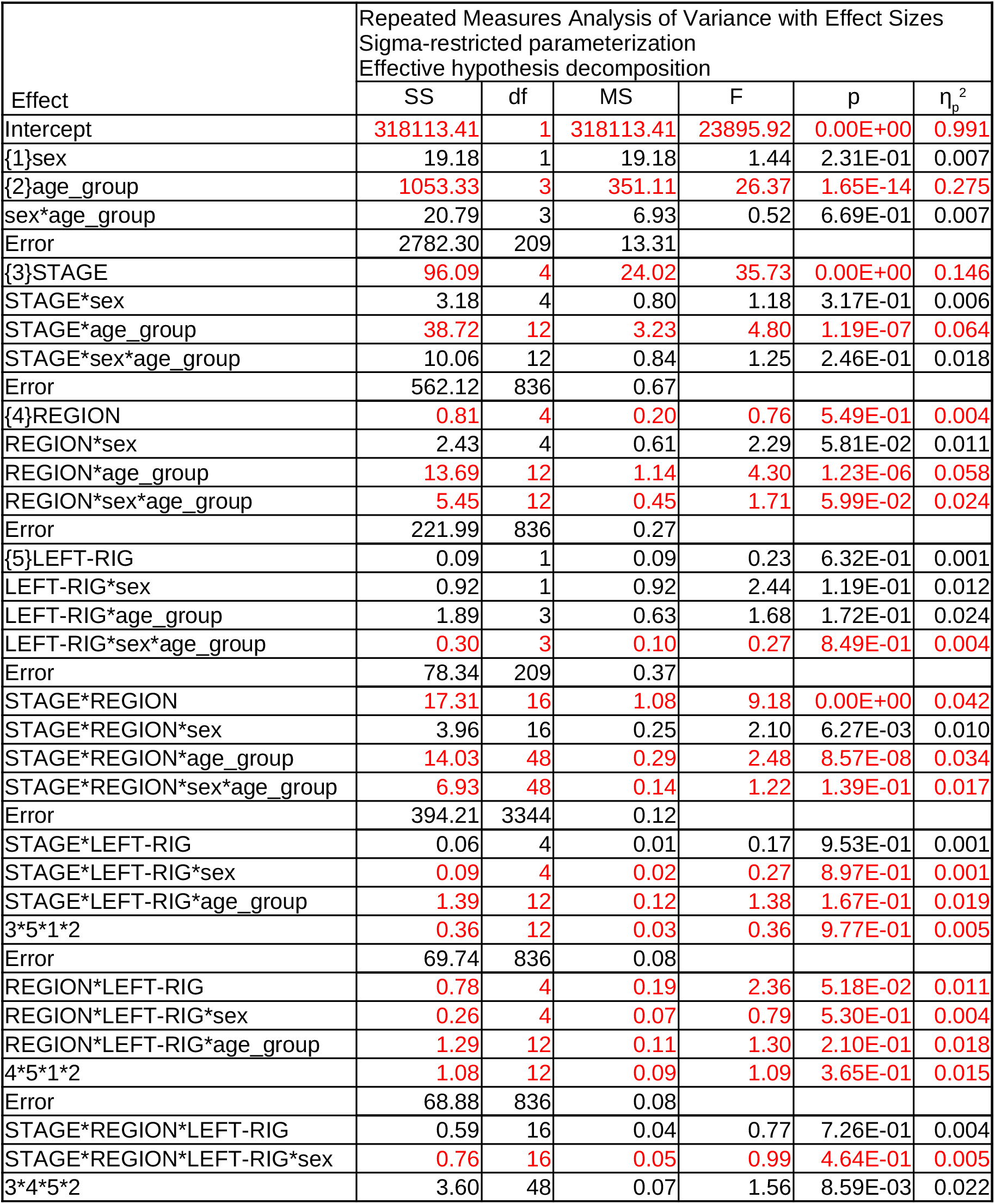

#### 7.4.3 Peak center frequency

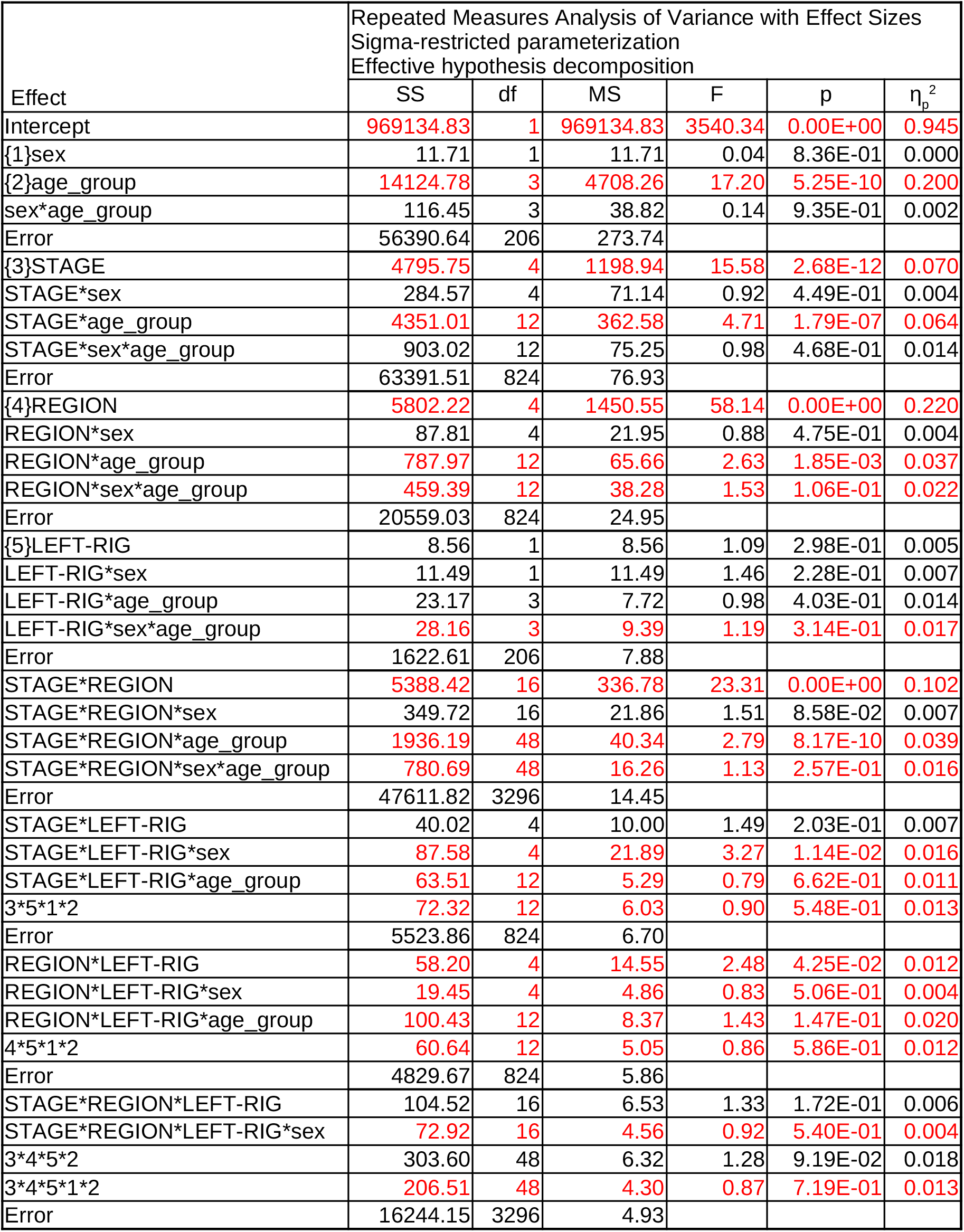

#### 7.4.4 Peak power

| Effect | Repeated Measures Analysis of Variance with Effect Size<br>Sigma-restricted parameterization<br>Effective hypothesis decomposition |  |  |  |  |  |
| --- | --- | --- | --- | --- | --- | --- |
| | SS | df | MS | F | p | $\eta_p^2$ |
| Intercept | 2909.38 | 1 | 2909.38 | 3215.24 | 0.00E+00 | 0.940 |
| {1}sex | 0.09 | 1 | 0.09 | 0.09 | 7.58E-01 | 0.000 |
| {2}age_group | 12.00 | 3 | 4.00 | 4.42 | 4.89E-03 | 0.060 |
| sex*age_group | 0.77 | 3 | 0.26 | 0.28 | 8.36E-01 | 0.004 |
| Error | 186.40 | 206 | 0.90 |  |  |  |
| {3}STAGE | 112.52 | 4 | 28.13 | 88.77 | 0.00E+00 | 0.301 |
| STAGE*sex | 0.64 | 4 | 0.16 | 0.50 | 7.34E-01 | 0.002 |
| STAGE*age_group | 18.58 | 12 | 1.55 | 4.88 | 8.01E-08 | 0.066 |
| STAGE*sex*age_group | 5.52 | 12 | 0.46 | 1.45 | 1.37E-01 | 0.021 |
| Error | 261.12 | 824 | 0.32 |  |  |  |
| {4}REGION | 19.08 | 4 | 4.77 | 97.65 | 0.00E+00 | 0.322 |
| REGION*sex | 0.32 | 4 | 0.08 | 1.64 | 1.61E-01 | 0.008 |
| REGION*age_group | 2.41 | 12 | 0.20 | 4.11 | 2.93E-06 | 0.056 |
| REGION*sex*age_group | 0.26 | 12 | 0.02 | 0.44 | 9.47E-01 | 0.006 |
| Error | 40.25 | 824 | 0.05 |  |  |  |
| {5}LEFT-RIG | 0.00 | 1 | 0.00 | 0.01 | 9.40E-01 | 0.000 |
| LEFT-RIG*sex | 0.01 | 1 | 0.01 | 0.68 | 4.09E-01 | 0.003 |
| LEFT-RIG*age_group | 0.05 | 3 | 0.02 | 1.37 | 2.52E-01 | 0.020 |
| LEFT-RIG*sex*age_group | 0.01 | 3 | 0.00 | 0.17 | 9.17E-01 | 0.002 |
| Error | 2.63 | 206 | 0.01 |  |  |  |
| STAGE*REGION | 14.20 | 16 | 0.89 | 47.27 | 0.00E+00 | 0.187 |
| STAGE*REGION*sex | 0.10 | 16 | 0.01 | 0.33 | 9.95E-01 | 0.002 |
| STAGE*REGION*age_group | 5.95 | 48 | 0.12 | 6.60 | 0.00E+00 | 0.088 |
| STAGE*REGION*sex*age_group | 1.01 | 48 | 0.02 | 1.12 | 2.58E-01 | 0.016 |
| Error | 61.90 | 3296 | 0.02 |  |  |  |
| STAGE*LEFT-RIG | 0.10 | 4 | 0.03 | 4.89 | 6.67E-04 | 0.023 |
| STAGE*LEFT-RIG*sex | 0.02 | 4 | 0.01 | 1.04 | 3.88E-01 | 0.005 |
| STAGE*LEFT-RIG*age_group | 0.17 | 12 | 0.01 | 2.71 | 1.31E-03 | 0.038 |
| 3*5*1*2 | 0.03 | 12 | 0.00 | 0.49 | 9.21E-01 | 0.007 |
| Error | 4.33 | 824 | 0.01 |  |  |  |
| REGION*LEFT-RIG | 0.04 | 4 | 0.01 | 1.18 | 3.18E-01 | 0.006 |
| REGION*LEFT-RIG*sex | 0.03 | 4 | 0.01 | 1.03 | 3.92E-01 | 0.005 |
| REGION*LEFT-RIG*age_group | 0.13 | 12 | 0.01 | 1.29 | 2.17E-01 | 0.018 |
| 4*5*1*2 | 0.11 | 12 | 0.01 | 1.08 | 3.72E-01 | 0.016 |
| Error | 6.92 | 824 | 0.01 |  |  |  |
| STAGE*REGION*LEFT-RIG | 0.13 | 16 | 0.01 | 2.46 | 9.93E-04 | 0.012 |
| STAGE*REGION*LEFT-RIG*sex | 0.07 | 16 | 0.00 | 1.29 | 1.91E-01 | 0.006 |
| 3*4*5*2 | 0.29 | 48 | 0.01 | 1.85 | 3.70E-04 | 0.026 |
| 3*4*5*1*2 | 0.22 | 48 | 0.00 | 1.38 | 4.48E-02 | 0.020 |
| Error | 10.82 | 3296 | 0.00 |  |  |  |

#### 7.4.5 Adjusted spectral slope

| Effect | Repeated Measures Analysis of Variance with Effect Sizes<br>Sigma-restricted parameterization<br>Effective hypothesis decomposition |  |  |  |  |  |
| --- | --- | --- | --- | --- | --- | --- |
| | SS | df | MS | F | p | $\eta_p^2$ |
| Intercept | 1688.622 | 1 | 1688.622 | 492.449 | 0.00E+00 | 0.702045 |
| {1}sex | 0.664 | 1 | 0.664 | 0.194 | 6.60E-01 | 0.000925 |
| {2}age_group | 36.531 | 3 | 12.177 | 3.551 | 1.53E-02 | 0.048501 |
| Error | 716.667 | 209 | 3.429 |  |  |  |
| {3}STAGE | 552.469 | 3 | 184.156 | 1198.557 | 0.00E+00 | 0.851516 |
| STAGE*sex | 0.240 | 3 | 0.080 | 0.520 | 6.68E-01 | 0.002484 |
| STAGE*age_group | 16.050 | 9 | 1.783 | 11.607 | 1.11E-16 | 0.142812 |
| Error | 96.338 | 627 | 0.154 |  |  |  |
| {4}REGION | 8.766 | 4 | 2.191 | 43.498 | 0.00E+00 | 0.172270 |
| REGION*sex | 0.321 | 4 | 0.080 | 1.592 | 1.74E-01 | 0.007559 |
| REGION*age_group | 1.485 | 12 | 0.124 | 2.456 | 3.76E-03 | 0.034051 |
| Error | 42.117 | 836 | 0.050 |  |  |  |
| {5}LEFT-RIG | 0.066 | 1 | 0.066 | 1.086 | 2.98E-01 | 0.005171 |
| LEFT-RIG*sex | 0.241 | 1 | 0.241 | 3.936 | 4.86E-02 | 0.018485 |
| LEFT-RIG*age_group | 0.411 | 3 | 0.137 | 2.241 | 8.45E-02 | 0.031172 |
| Error | 12.782 | 209 | 0.061 |  |  |  |
| STAGE*REGION | 3.834 | 12 | 0.319 | 62.999 | 0.00E+00 | 0.231614 |
| STAGE*REGION*sex | 0.080 | 12 | 0.007 | 1.316 | 2.02E-01 | 0.006255 |
| STAGE*REGION*age_group | 1.201 | 36 | 0.033 | 6.577 | 0.00E+00 | 0.086263 |
| Error | 12.718 | 2508 | 0.005 |  |  |  |
| STAGE*LEFT-RIG | 0.027 | 3 | 0.009 | 1.295 | 2.75E-01 | 0.006156 |
| STAGE*LEFT-RIG*sex | 0.005 | 3 | 0.002 | 0.260 | 8.54E-01 | 0.001241 |
| STAGE*LEFT-RIG*age_group | 0.055 | 9 | 0.006 | 0.869 | 5.52E-01 | 0.012326 |
| Error | 4.402 | 627 | 0.007 |  |  |  |
| REGION*LEFT-RIG | 0.098 | 4 | 0.024 | 1.729 | 1.42E-01 | 0.008204 |
| REGION*LEFT-RIG*sex | 0.013 | 4 | 0.003 | 0.236 | 9.18E-01 | 0.001128 |
| REGION*LEFT-RIG*age_group | 0.129 | 12 | 0.011 | 0.762 | 6.90E-01 | 0.010826 |
| Error | 11.830 | 836 | 0.014 |  |  |  |
| STAGE*REGION*LEFT-RIG | 0.010 | 12 | 0.001 | 0.621 | 8.26E-01 | 0.002964 |
| STAGE*REGION*LEFT-RIG*sex | 0.015 | 12 | 0.001 | 0.967 | 4.79E-01 | 0.004603 |
| 3*4*5*2 | 0.046 | 36 | 0.001 | 0.997 | 4.75E-01 | 0.014104 |
| Error | 3.220 | 2508 | 0.001 |  |  |  |

