## Supplementary material for "Scale-free and oscillatory spectral measures of sleep stages in humans": Slope statistics (from another version of this preprint: version 1, 2022-06-12)

| Effect | Repeated Measures Analysis of Variance with Effect Sizes<br>Sigma-restricted parameterization<br>Effective hypothesis decomposition | | | | | |
|---|---|---|---|---|---|---|
| | SS | df | MS | F | p | $\eta_p^2$ |
| Intercept | 37765.20 | 1 | 37765.20 | 30064.52 | 0.00E+00 | 0.993 |
| {1}sex | 0.18 | 1 | 0.18 | 0.15 | 7.02E-01 | 0.001 |
| {2}age_group | 24.38 | 3 | 8.13 | 6.47 | 3.32E-04 | 0.086 |
| sex*age_group | 3.00 | 3 | 1.00 | 0.80 | 4.97E-01 | 0.011 |
| Error | 258.76 | 206 | 1.26 | | | |
| {3}STAGE | 882.03 | 4 | 220.51 | 770.29 | 0.00E+00 | 0.789 |
| STAGE*sex | 1.27 | 4 | 0.32 | 1.11 | 3.49E-01 | 0.005 |
| STAGE*age_group | 23.71 | 12 | 1.98 | 6.90 | 5.37E-12 | 0.091 |
| STAGE*sex*age_group | 3.79 | 12 | 0.32 | 1.10 | 3.54E-01 | 0.016 |
| Error | 235.88 | 824 | 0.29 | | | |
| {4}REGION | 22.42 | 4 | 5.60 | 113.33 | 0.00E+00 | 0.355 |
| REGION*sex | 0.36 | 4 | 0.09 | 1.83 | 1.21E-01 | 0.009 |
| REGION*age_group | 3.86 | 12 | 0.32 | 6.51 | 3.50E-11 | 0.087 |
| REGION*sex*age_group | 1.05 | 12 | 0.09 | 1.76 | 4.98E-02 | 0.025 |
| Error | 40.75 | 824 | 0.05 | | | |
| {5}LEFT-RIG | 0.08 | 1 | 0.08 | 2.56 | 1.11E-01 | 0.012 |
| LEFT-RIG*sex | 0.07 | 1 | 0.07 | 2.13 | 1.46E-01 | 0.010 |
| LEFT-RIG*age_group | 0.17 | 3 | 0.06 | 1.82 | 1.45E-01 | 0.026 |
| LEFT-RIG*sex*age_group | 0.27 | 3 | 0.09 | 2.95 | 3.39E-02 | 0.041 |
| Error | 6.32 | 206 | 0.03 | | | |
| STAGE*REGION | 5.54 | 16 | 0.35 | 55.23 | 0.00E+00 | 0.211 |
| STAGE*REGION*sex | 0.13 | 16 | 0.01 | 1.27 | 2.10E-01 | 0.006 |
| STAGE*REGION*age_group | 1.49 | 48 | 0.03 | 4.95 | 0.00E+00 | 0.067 |
| STAGE*REGION*sex*age_group | 0.47 | 48 | 0.01 | 1.55 | 9.54E-03 | 0.022 |
| Error | 20.68 | 3296 | 0.01 | | | |
| STAGE*LEFT-RIG | 0.05 | 4 | 0.01 | 1.52 | 1.93E-01 | 0.007 |
| STAGE*LEFT-RIG*sex | 0.11 | 4 | 0.03 | 3.42 | 8.82E-03 | 0.016 |
| STAGE*LEFT-RIG*age_group | 0.18 | 12 | 0.02 | 1.87 | 3.41E-02 | 0.027 |
| 3*5*1*2 | 0.30 | 12 | 0.02 | 3.09 | 2.75E-04 | 0.043 |
| Error | 6.66 | 824 | 0.01 | | | |
| REGION*LEFT-RIG | 0.49 | 4 | 0.12 | 10.09 | 5.60E-08 | 0.047 |
| REGION*LEFT-RIG*sex | 0.20 | 4 | 0.05 | 4.07 | 2.82E-03 | 0.019 |
| REGION*LEFT-RIG*age_group | 0.40 | 12 | 0.03 | 2.75 | 1.13E-03 | 0.039 |
| 4*5*1*2 | 0.38 | 12 | 0.03 | 2.65 | 1.75E-03 | 0.037 |
| Error | 9.95 | 824 | 0.01 | | | |
| STAGE*REGION*LEFT-RIG | 0.03 | 16 | 0.00 | 1.27 | 2.04E-01 | 0.006 |
| STAGE*REGION*LEFT-RIG*sex | 0.04 | 16 | 0.00 | 1.70 | 3.96E-02 | 0.008 |
| 3*4*5*2 | 0.08 | 48 | 0.00 | 0.98 | 5.08E-01 | 0.014 |
| 3*4*5*1*2 | 0.17 | 48 | 0.00 | 2.11 | 1.48E-05 | 0.030 |
| Error | 5.42 | 3296 | 0.00 | | | |
