## Supplementary material for "Scale-free and oscillatory spectral measures of sleep stages in humans": Central frequency statistics (from another version of this preprint: version 1, 2022-06-12)

| Effect | Repeated Measures Analysis of Variance with Effect Sizes<br>Sigma-restricted parameterization<br>Effective hypothesis decomposition | | | | | |
|---|---|---|---|---|---|---|
| | SS | df | MS | F | p | $\eta_p^2$ |
| Intercept | 969134.83 | 1 | 969134.83 | 3540.34 | 0.00E+00 | 0.945 |
| {1}sex | 11.71 | 1 | 11.71 | 0.04 | 8.36E-01 | 0.000 |
| {2}age_group | 14124.78 | 3 | 4708.26 | 17.20 | 5.25E-10 | 0.200 |
| sex*age_group | 116.45 | 3 | 38.82 | 0.14 | 9.35E-01 | 0.002 |
| Error | 56390.64 | 206 | 273.74 | | | |
| {3}STAGE | 4795.75 | 4 | 1198.94 | 15.58 | 2.68E-12 | 0.070 |
| STAGE*sex | 284.57 | 4 | 71.14 | 0.92 | 4.49E-01 | 0.004 |
| STAGE*age_group | 4351.01 | 12 | 362.58 | 4.71 | 1.79E-07 | 0.064 |
| STAGE*sex*age_group | 903.02 | 12 | 75.25 | 0.98 | 4.68E-01 | 0.014 |
| Error | 63391.51 | 824 | 76.93 | | | |
| {4}REGION | 5802.22 | 4 | 1450.55 | 58.14 | 0.00E+00 | 0.220 |
| REGION*sex | 87.81 | 4 | 21.95 | 0.88 | 4.75E-01 | 0.004 |
| REGION*age_group | 787.97 | 12 | 65.66 | 2.63 | 1.85E-03 | 0.037 |
| REGION*sex*age_group | 459.39 | 12 | 38.28 | 1.53 | 1.06E-01 | 0.022 |
| Error | 20559.03 | 824 | 24.95 | | | |
| {5}LEFT-RIG | 8.56 | 1 | 8.56 | 1.09 | 2.98E-01 | 0.005 |
| LEFT-RIG*sex | 11.49 | 1 | 11.49 | 1.46 | 2.28E-01 | 0.007 |
| LEFT-RIG*age_group | 23.17 | 3 | 7.72 | 0.98 | 4.03E-01 | 0.014 |
| LEFT-RIG*sex*age_group | 28.16 | 3 | 9.39 | 1.19 | 3.14E-01 | 0.017 |
| Error | 1622.61 | 206 | 7.88 | | | |
| STAGE*REGION | 5388.42 | 16 | 336.78 | 23.31 | 0.00E+00 | 0.102 |
| STAGE*REGION*sex | 349.72 | 16 | 21.86 | 1.51 | 8.58E-02 | 0.007 |
| STAGE*REGION*age_group | 1936.19 | 48 | 40.34 | 2.79 | 8.17E-10 | 0.039 |
| STAGE*REGION*sex*age_group | 780.69 | 48 | 16.26 | 1.13 | 2.57E-01 | 0.016 |
| Error | 47611.82 | 3296 | 14.45 | | | |
| STAGE*LEFT-RIG | 40.02 | 4 | 10.00 | 1.49 | 2.03E-01 | 0.007 |
| STAGE*LEFT-RIG*sex | 87.58 | 4 | 21.89 | 3.27 | 1.14E-02 | 0.016 |
| STAGE*LEFT-RIG*age_group | 63.51 | 12 | 5.29 | 0.79 | 6.62E-01 | 0.011 |
| 3*5*1*2 | 72.32 | 12 | 6.03 | 0.90 | 5.48E-01 | 0.013 |
| Error | 5523.86 | 824 | 6.70 | | | |
| REGION*LEFT-RIG | 58.20 | 4 | 14.55 | 2.48 | 4.25E-02 | 0.012 |
| REGION*LEFT-RIG*sex | 19.45 | 4 | 4.86 | 0.83 | 5.06E-01 | 0.004 |
| REGION*LEFT-RIG*age_group | 100.43 | 12 | 8.37 | 1.43 | 1.47E-01 | 0.020 |
| 4*5*1*2 | 60.64 | 12 | 5.05 | 0.86 | 5.86E-01 | 0.012 |
| Error | 4829.67 | 824 | 5.86 | | | |
| STAGE*REGION*LEFT-RIG | 104.52 | 16 | 6.53 | 1.33 | 1.72E-01 | 0.006 |
| STAGE*REGION*LEFT-RIG*sex | 72.92 | 16 | 4.56 | 0.92 | 5.40E-01 | 0.004 |
| 3*4*5*2 | 303.60 | 48 | 6.32 | 1.28 | 9.19E-02 | 0.018 |
| 3*4*5*1*2 | 206.51 | 48 | 4.30 | 0.87 | 7.19E-01 | 0.013 |
| Error | 16244.15 | 3296 | 4.93 | | | |
