## Supplementary material for "Scale-free and oscillatory spectral measures of sleep stages in humans": Intercept statistics (from another version of this preprint: version 1, 2022-06-12)

| Effect | Repeated Measures Analysis of Variance with Effect Sizes<br>Sigma-restricted parameterization<br>Effective hypothesis decomposition | | | | | |
|---|---|---|---|---|---|---|
| | SS | df | MS | F | p | $\eta_p^2$ |
| Intercept | 318113.41 | 1 | 318113.41 | 23895.92 | 0.00E+00 | 0.991 |
| {1}sex | 19.18 | 1 | 19.18 | 1.44 | 2.31E-01 | 0.007 |
| {2}age_group | 1053.33 | 3 | 351.11 | 26.37 | 1.65E-14 | 0.275 |
| sex*age_group | 20.79 | 3 | 6.93 | 0.52 | 6.69E-01 | 0.007 |
| Error | 2782.30 | 209 | 13.31 | | | |
| {3}STAGE | 96.09 | 4 | 24.02 | 35.73 | 0.00E+00 | 0.146 |
| STAGE*sex | 3.18 | 4 | 0.80 | 1.18 | 3.17E-01 | 0.006 |
| STAGE*age_group | 38.72 | 12 | 3.23 | 4.80 | 1.19E-07 | 0.064 |
| STAGE*sex*age_group | 10.06 | 12 | 0.84 | 1.25 | 2.46E-01 | 0.018 |
| Error | 562.12 | 836 | 0.67 | | | |
| {4}REGION | 0.81 | 4 | 0.20 | 0.76 | 5.49E-01 | 0.004 |
| REGION*sex | 2.43 | 4 | 0.61 | 2.29 | 5.81E-02 | 0.011 |
| REGION*age_group | 13.69 | 12 | 1.14 | 4.30 | 1.23E-06 | 0.058 |
| REGION*sex*age_group | 5.45 | 12 | 0.45 | 1.71 | 5.99E-02 | 0.024 |
| Error | 221.99 | 836 | 0.27 | | | |
| {5}LEFT-RIG | 0.09 | 1 | 0.09 | 0.23 | 6.32E-01 | 0.001 |
| LEFT-RIG*sex | 0.92 | 1 | 0.92 | 2.44 | 1.19E-01 | 0.012 |
| LEFT-RIG*age_group | 1.89 | 3 | 0.63 | 1.68 | 1.72E-01 | 0.024 |
| LEFT-RIG*sex*age_group | 0.30 | 3 | 0.10 | 0.27 | 8.49E-01 | 0.004 |
| Error | 78.34 | 209 | 0.37 | | | |
| STAGE*REGION | 17.31 | 16 | 1.08 | 9.18 | 0.00E+00 | 0.042 |
| STAGE*REGION*sex | 3.96 | 16 | 0.25 | 2.10 | 6.27E-03 | 0.010 |
| STAGE*REGION*age_group | 14.03 | 48 | 0.29 | 2.48 | 8.57E-08 | 0.034 |
| STAGE*REGION*sex*age_group | 6.93 | 48 | 0.14 | 1.22 | 1.39E-01 | 0.017 |
| Error | 394.21 | 3344 | 0.12 | | | |
| STAGE*LEFT-RIG | 0.06 | 4 | 0.01 | 0.17 | 9.53E-01 | 0.001 |
| STAGE*LEFT-RIG*sex | 0.09 | 4 | 0.02 | 0.27 | 8.97E-01 | 0.001 |
| STAGE*LEFT-RIG*age_group | 1.39 | 12 | 0.12 | 1.38 | 1.67E-01 | 0.019 |
| 3*5*1*2 | 0.36 | 12 | 0.03 | 0.36 | 9.77E-01 | 0.005 |
| Error | 69.74 | 836 | 0.08 | | | |
| REGION*LEFT-RIG | 0.78 | 4 | 0.19 | 2.36 | 5.18E-02 | 0.011 |
| REGION*LEFT-RIG*sex | 0.26 | 4 | 0.07 | 0.79 | 5.30E-01 | 0.004 |
| REGION*LEFT-RIG*age_group | 1.29 | 12 | 0.11 | 1.30 | 2.10E-01 | 0.018 |
| 4*5*1*2 | 1.08 | 12 | 0.09 | 1.09 | 3.65E-01 | 0.015 |
| Error | 68.88 | 836 | 0.08 | | | |
| STAGE*REGION*LEFT-RIG | 0.59 | 16 | 0.04 | 0.77 | 7.26E-01 | 0.004 |
| STAGE*REGION*LEFT-RIG*sex | 0.76 | 16 | 0.05 | 0.99 | 4.64E-01 | 0.005 |
| 3*4*5*2 | 3.60 | 48 | 0.07 | 1.56 | 8.59E-03 | 0.022 |

| | | | | | | |
|---|---|---|---|---|---|---|
| 3*4*5*1*2 | 2.53 | 48 | 0.05 | 1.09 | 3.04E-01 | 0.015 |
| Error | 160.97 | 3344 | 0.05 | | | |
